# Tannic Acid Inhibits α-Synuclein Amyloid Fibril Formation via Binding to the Monomer N-terminal Domain

**DOI:** 10.1101/2021.05.26.445860

**Authors:** Jonathan Stoeber, Jonathan K. Williams, Prabhas V. Moghe, Jean Baum

## Abstract

α-Synuclein (αS) is an intrinsically disordered protein (IDP) that aggregates into amyloid fibrils during the progression of Parkinson’s Disease and other synucleinopathies. The N-terminal domain (residues 1-60) is now understood to play a critical role in the initial nucleation of aggregation, as well as a pivotal role in the monomer-fibril interaction underlying amyloid seeding. Here we report on the interaction between αS and the polyphenol tannic acid (TA), where a combination of solution NMR, atomic force microscopy (AFM), and ThT assays have identified that TA targets the αS N-terminal domain to inhibit amyloid fibril formation in a pH dependent manner. This work highlights the importance of targeting the N-terminus of αS to arrest fibril formation, and suggests the importance of including polyphenolic moieties in future amyloid inhibitors.

## 1. Introduction

α-synuclein (αS) is an intrinsically disordered protein (IDP) that is implicated in the pathogenesis of Parkinson’s Disease (PD). The structural plasticity and disordered nature of the αS IDP monomer is believed to aid its function *in vivo* [1], evident by its adoption of an α-helical structure when membrane bound [2]. However, there are disagreements as to whether αS forming α-helices inhibits aggregation [3,4] or induces fibril formation [5]. In PD, αS misfolds and undergoes a structural transition into β-sheet containing higher ordered oligomers [6] and fibrils [7]. αS has three distinct domains: the negatively charged, amphipathic N-terminus (residues 1-60) containing four imperfect KTKEGV motifs, responsible for membrane binding but also might cause aggregation [8,9]; the hydrophobic, non-amyloid component (NAC) domain (residues 61-95) which makes up the amyloid fibril core and is necessary for fibril formation [10,11]; and lastly the highly acidic and proline rich C-terminus (residues 96-140), which has been shown to interact with calcium [12], chaperones [13], and other IDPs [14–16]. Gaining a deeper understanding of how αS interacts with its various binding partners will not only shed light on the factors that define its normal function and drive the earliest stages of its misfolding and aggregation, but also provide fundamental insight into αS-ligand interactions, which can then be exploited to prevent the disordered-to-ordered structural transition leading to disease related aggregation.

Polyphenols are a class of compounds that contain numerous aromatic groups that have vicinal hydroxyl moieties, and have been shown to inhibit amyloid fibril formation and disaggregate pre-formed fibrils [17–24]. For example, the compounds brazilin and epigallocatechin-3-gallate (EGCG) were found to inhibit the aggregation of amyloid-β (Aβ) and αS, respectively, through the formation of off-pathway aggregates [25–27]. However, how these polyphenols inhibit aggregation is not well understood and can differ depending on the specific polyphenol, including how it interacts with the protein and at what stage of aggregation it stabilizes [19]. Determining how different polyphenolic compounds inhibit IDP aggregation and how this relates to the structure of the compound will aid in understanding the mechanisms of IDP aggregation in addition to informing the design of novel polyphenolic therapeutics. The branched polyphenolic compound, Tannic acid (TA), has presented itself as a promising candidate for inhibiting the aggregation of αS. Preliminary studies have shown that TA, similar to EGCG and brazilin, inhibits the aggregation and destabilizes preformed fibrils of both αS and Aβ [17,28]. It was also shown to be the most potent inhibitor of αS aggregation among a group of 11 different antioxidants [29]. Our previous work reported that TA within nanoparticles attenuated αS fibrillization and intracellular aggregation within microglia, with beneficial effects on neuroinflammatory pathways [30]. However, it is still not understood how TA interacts with αS mechanistically and inhibits its aggregation.

In this work, we have determined the residue-specific interactions between monomeric αS and TA using solution NMR spectroscopy, along with ThT aggregation assays and AFM imaging, to understand how TA interacts with and affects αS aggregation. We observed that TA’s ability to interact with αS is pH dependent, likely through autooxidation, which promotes TA interaction with αS and inhibits fibril formation at neutral pH. Thioflavin T (ThT) fluorescence assays showed that TA inhibits αS in a concentration dependent manner, with full inhibition observed at a 1:1 αS:TA molar ratio. A combination of atomic force microscopy (AFM), size-exclusion chromatography (SEC), and sodium dodecyl sulphate polyacrylamide gel electrophoresis (SDS-PAGE) showed that TA induces the formation of oligomers of various sizes, both soluble and insoluble. Solution NMR ^1^H-^15^N heteronuclear single quantum coherence (HSQC) and ^13^C-^15^N carbonyl-nitrogen (CON) correlation spectra, along with residue specific ^15^N-R_2_ relaxation rates, indicated that TA primarily interacts with the N-terminal domain of αS or with positively charged or polar residues in the other domains. Our work here highlights the importance of targeting the N-terminal domain for effective inhibition of αS fibril formation, and indicate that oxidized derivatives of branched-polyphenolic compounds may be interesting targets for future development of amyloid inhibitors.

## 2. Results

### 2.1 TA Becomes an Efficient Inhibitor of αS Amyloid Fibril Formation at pH 7.4

To determine the effects of TA on the kinetics of αS aggregation, ThT assays were conducted at various αS:TA molar ratios at both pH 7.4 (Fig. 1a) and pH 6.0 (Fig. S2a). ThT is a amyloid-active fluorescent dye that is well established and widely used to study amyloid protein aggregation [31]. We observed a concentration dependent inhibition of αS aggregation with increasing concentrations of TA, with significant inhibition of fibril formation occurring at aS:TA molar ratios of 10:1 (Fig. 1a, green) and complete inhibition at a 1:1 αS:TA ratio (Fig. 1a, orange). At pH 6.0 no significant inhibition of αS aggregation by TA was observed at any ratio of aS:TA (Fig. S2a). Interestingly at pH 6.0 TA exhibits some potential acceleration of αS aggregation at a 1:1 aS:TA ratio (Fig. S2a, orange). This behavior is similar to previous observations of the interaction between the polyphenol EGCG and αS [32,33]. To ensure that TA does not interfere with the ability of ThT to fluoresce upon binding to amyloid fibrils and to gain a detailed picture of the aggregates formed in the presence of TA, we acquired AFM images of the endpoints of the αS control (Fig. 1b) and the 1:1 αS:TA (Fig. 1c) assays at pH 7.4. These AFM images show that TA almost completely prevents the formation αS fibrils, and instead results in the formation of small amorphous aggregate species. To assess the amount of these αS species that are present in the soluble fraction at the various molar ratios of aS:TA used in the ThT assays, αS was incubated with various concentrations of TA for 7 days. The solution was then spun down and the supernatant (i.e. soluble fraction) was subjected to analysis by SDS-PAGE (Fig. S4). The results show that TA does not decrease the level of αS in the soluble phase until a αS:TA ratio of 1:1 is reached, the ratio of TA that completely inhibits fibril formation. The size of the species present in the soluble fraction was then analyzed via size-exclusion chromatography (SEC) (Fig. S5). The SEC profile shows 4 peaks in the 8-13 mL elution volume range, indicating that an assortment of soluble aggregate species are present. This indicates that at a molar ratio of 1:1 aS:TA, TA promotes the formation of soluble aggregate species instead of insoluble amyloid fibrils.

**Figure 1.**
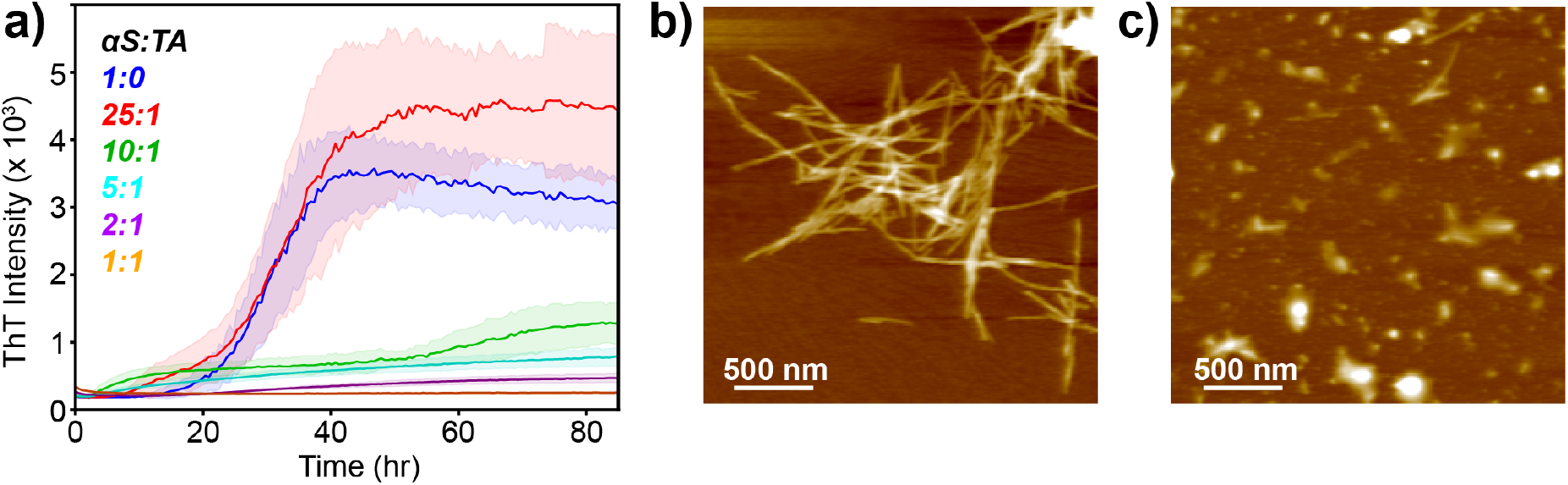
Tannic acid modulates the kinetics and final products formed during αS aggregation. (a) ThT fluorescence assays of αS fibril formation with increasing concentrations of TA: control 1:0 (blue), 25:1 (red), 10:1 (green), 5:1 (cyan), 2:1 (purple), 1:1 (orange). All assays were conducted in 100 mM PBS buffer at pH 7.4 with an αS concentration of 70 μM, 37°C and 600 rpm shaking. (b) AFM image of the fibrils formed from the αS control (blue curve) ThT assay, taken from the endpoint. (c) AFM image of the aggregates formed frm the αS:TA 1:1 (orange curve) ThT assay, taken from the endpoint.

### 2.2 Tannic Acid Likely Autooxidizes at pH 7.4

Because of the propensity for catechol compounds to autoxidize at neutral pH, we sought to gain an understanding of the oxidation state of TA in solution at both pH 7.4 and pH 6.0 using a combination of UV-Vis spectroscopy and reverse-phase high-pressure liquid chromatography (HPLC). As observed in the UV-Vis spectrum (Fig. S1a), TA has an absorbance maximum at 275 nm at pH 6.0, in good agreement with the literature (Absmax = 280 nm at pH 4.7) [34]. This maximum absorbance wavelength remains after the pH is increased to 7.4, but broadens within the range of 275-325 nm. In contrast, GA displays an absorbance maximum around 260 nm (Fig. S1b), in agreement with the literature (Absmax = 260 nm at pH 4.7), which remains relatively unchanged as the pH is increased from 6.0 to 7.4 [34]. No visible color change is detected upon increasing the pH of TA solutions from ~4 or 5 to pH 6.0, while the color of TA solutions in PBS changes from a pale orange to a dark brown once the pH reaches ~7.0. This pH induced color shift is consistent with the changes in the UV-Vis spectrum and HPLC trace (Fig. Slc) of TA at pH 7.4 relative to pH 6.0. Because of these changes, it is likely that TA is undergoing pH induced autooxidation at pH 7.4 where the vicinal hydroxyl groups form an assortment of branched ortho-quinone (o-quinone) like compounds [32,33,35].

### 2.3 Solution NMR Indicates that TA Interacts with the N-terminal Domain of αS

Solution NMR is a powerful technique that provides atomic-resolution and residue-level information on protein structure and dynamics, and can probe the ligand-induced changes to the local chemical environment of a protein. Adding a ligand to a protein solution can cause changes in the chemical shifts of a peak, called chemical shift perturbation (CSP), which indicate that there are interactions between the ligand and that residue. In addition to CSP, a loss of peak intensity or broadening of a peak is also indicative of interactions between protein and ligand, due to changes in protein dynamics from ligand binding. Observing these changes in NMR spectra of IDPs is complicated because of small chemical shift dispersions inherent to the disordered and unstructured nature of the IDPs.

To overcome issues of peak overlap inherent to the narrow ^1^H chemical shift dispersion of NMR spectra of IDPs, we utilized a combination of ^1^H-^15^N HSQC (Fig. 2) and ^15^N-^13^C CON (Fig. 3) correlation spectra to identify the residue-specific interactions between αS and TA at pH 7.4. The ^1^H, ^13^C, and ^15^N chemical shifts of αS at pH 7.4 have been well documented in the literature [36,37]. Based on these assignments, we observed that TA induces either chemical shift perturbation (CSP) or loss of peak intensity in 51 N-terminal residues, 18 NAC residues, and 8 C-terminal residues in either the HSQC or CON spectra (Fig. S6). The ^1^H-^15^N HSQC and ^15^N-^13^C CON spectra indicate that TA interacts primarily with the N-terminus, and with regions of the NAC domain that contain threonine residues. The ^15^N-^13^C CON spectrum in particular allowed for the detection of αS/TA interactions in the C-terminus (L100-G101, S129-E130) that were overlapped in the HSQC spectrum. At pH 6.0 the ^1^H-^15^N HSQC spectrum (Fig. S3) displays no CSP or peak intensity loss, suggesting that it is necessary for TA to undergo the autooxidation to a branched ortho-quinone compound for it to interact with αS.

**Figure 2.**
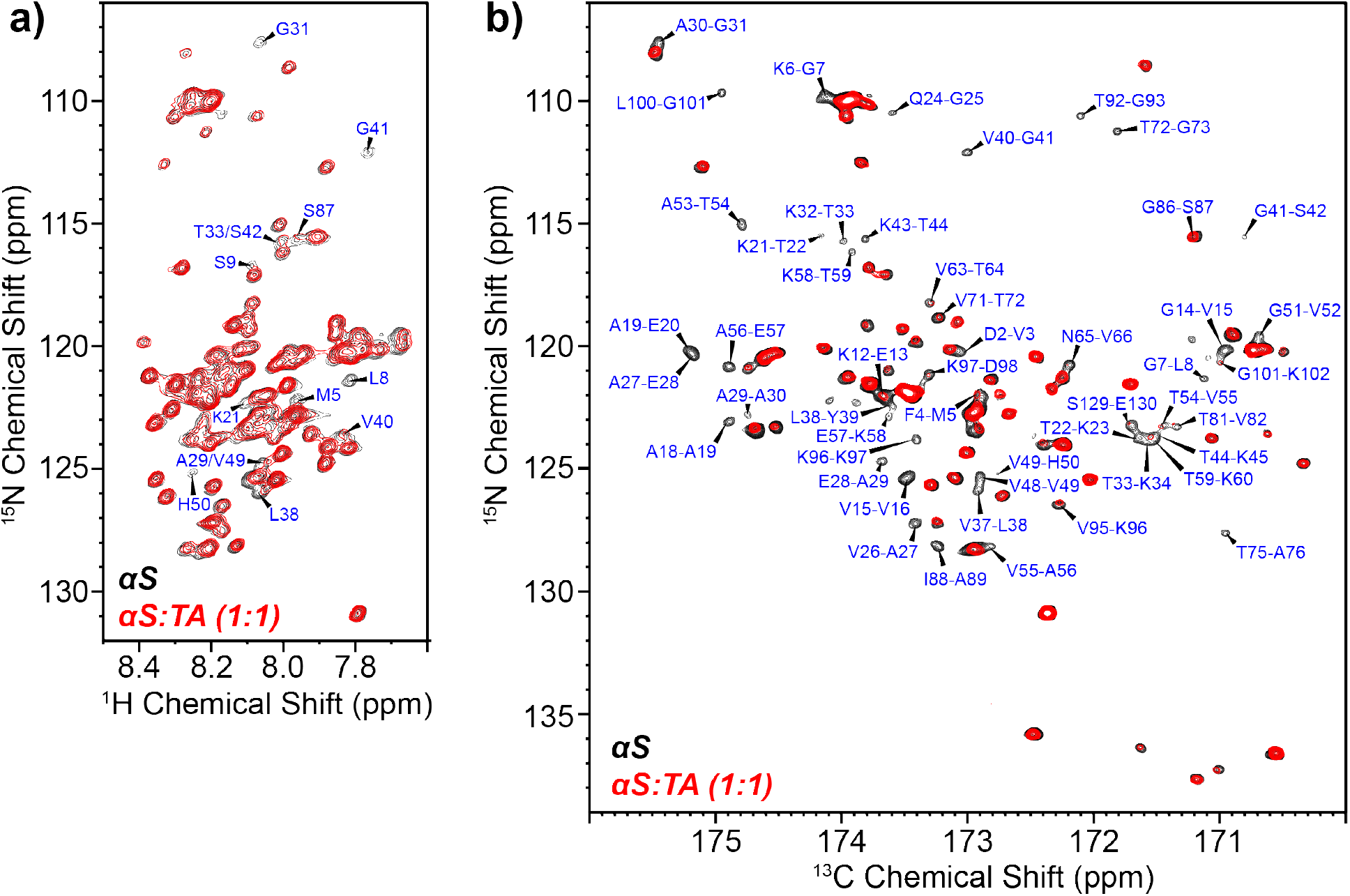
Tannic acid interacts with monomeric αS residues primarily located in the N-terminal domain. (a) ^1^H-^15^N HSQC spectra of monomeric αS (black) and αS:TA at a molar ratio of 1:1 (red). Many residues have cross peaks that are overlapped in this type of spectrum. The cross peaks that show reduced intensity or chemical shift perturbation in the presence of TA are labeled in blue. (b) ^15^N-^13^C CON spectra of monomeric αS (black) and αS:TA at a molar ratio of 1:1 (red). The majority of cross peaks in this type of spectrum are well resolved and do not overlap with one another, allowing for easy assignment. The cross peaks that show reduced intensity or chemical shift perturbation in the presence of TA are labeled in blue.

**Figure 3.**
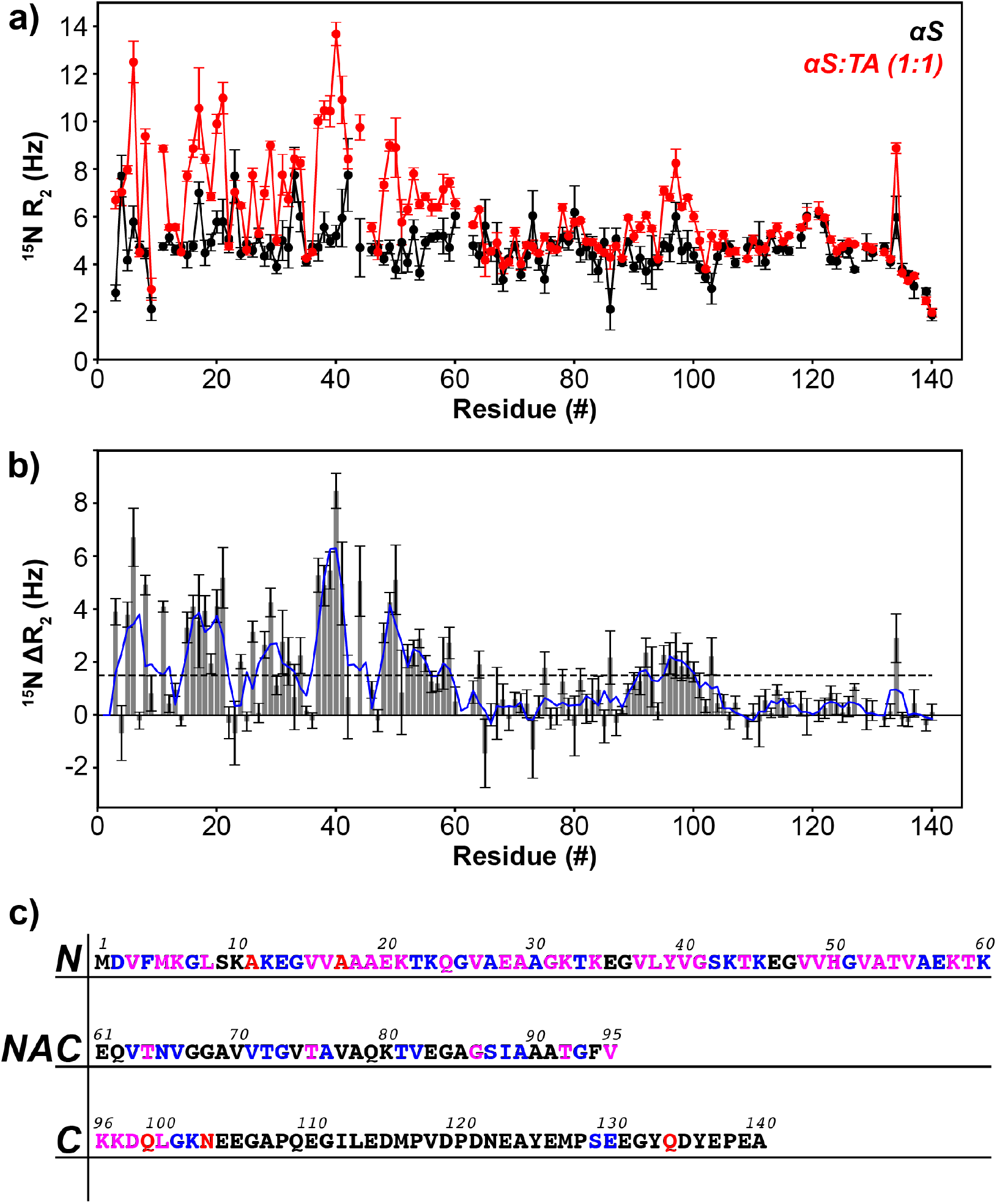
Tannic acid increases ^15^N-R_2_ relaxation rates primarily in the N-terminal domain. (a) Residue-specific ^15^N-R_2_ relaxation rates of monomeric αS (black) and αS:TA at molar ratio of 1:1 (red). (b) The difference in ^15^N-R_2_ relaxation rates between αS:TA and αS (ΔR_2_) along the αS monomer. A running average of every 3 residues is plotted as a blue line. A dashed grey line is included to demarcate a ΔR_2_ of 1.5 Hz. (c) The αS sequence is shown for each of its three domains: N-terminal (1-60), NAC (61-95), C-terminal (96-140). Residues that were found to have chemical shift perturbations or loss of intensity in the HSQC and CON spectra are shown in blue, residues that had increases in ^15^N-R_2_ greater than 1.5 Hz are shown in red, and residues that showed both a CSP/intensity loss and an increase in R_2_ are shown in magenta. Residues that did not meet any of these criteria are in black.

To further probe the interactions between αS and TA, ^15^N-R_2_ relaxation NMR experiments were conducted (Fig. 4) before and after the addition of TA in order to correlate the residue-specific changes in backbone dynamics induced via interactions with the ligand [38,39]. ^15^N-R_2_ relaxation experiments were conducted for αS and a 1:1 ratio of αS:TA (Fig. 1a). Residues with a significant change in ^15^N-R_2_ (ΔR_2_ > 1.5 Hz, Fig. 3b) are shown in Figure 3c, highlighted in red. The majority of residues displaying a significant change in R_2_ are found in the N-terminus, in agreement to the peak intensity losses observed in the CON spectra. Residues that show significant changes in ^15^N-R_2_ in the NAC (T64, T75, G86, T92, and V95) and C-terminal domains (K96, K97, D98, Q99, L100, N103, and Q134) are in the same regions that also displayed reduced peak intensity in the CON spectra. The changes in ^15^N-R_2_ values observed at a 1:1 ratio of αS:TA indicate that TA interacts primarily with the N-terminus of αS, and a few other polar or positively charged regions of the NAC region and C-terminal domains.

## 3. Discussion

Identifying a small molecule that can prevent αS fibril formation, or interrupt fibril seeding and propagation, is of intense interest to the treatment of Parkinson’s disease and other synucleinopathies. Many compounds have been found to modulate the aggregation behavior of αS, and some have even been found to disaggregate already formed oligomers and fibrils [17,21,40]. Several chemical moieties have been identified that show particularly strong effects for inhibiting αS fibril formation, including polyphenols, rifamycins, terpenoids, and phenothiazines [41]. The polyphenols in particular have been studied in depth, due to their natural occurrence and potential health benefits. Tannic acid (TA) and epigallocatechin gallate (EGCG) are polyphenolic compounds which have previously been found to inhibit αS aggregation [17,27]. However, the conditions under which these compounds are able to successfully prevent amyloid aggregation are varied. EGCG was found to inhibit insulin fibril formation at pH 7.0 but not at pH 6.0 [32], while it completely inhibits αS aggregation at pH 7.0 and actually accelerates aggregation at pH 6.0 [33]. The results of the current work show that TA is an efficient inhibitor of αS at pH 7.4 but not at pH 6.0, which is consistent with previous work in the literature utilizing similar polyphenols and IDPs [32,33]. The pH dependent inhibitory behavior of these polyphenols is thought to be a result of autooxidation at neutral pH [42,43]. Under these conditions the vicinal hydroxyl groups of the catechol moieties of the polyphenols can form o-quinone intermediates that have the potential to form conjugates with the sidechain amino group of lysine and guanidinium group of arginine [23,40]. This is in agreement with our observations here, where TA has induced loss of cross peak intensity and increases in ^15^N-R_2_ for the majority of the αS N-terminal domain, which is where most of the lysine residues are located in KTKEGV repeats, as well as the very beginning of the C-terminal domain where 3 more lysine residues reside. Previous work has determined that vicinal hydroxyl moieties were necessary for flavonoids to be amyloid inhibitors, and a third vicinal hydroxyl group increased inhibitory properties [23,35]. The exact stoichiometry of the TA-αS interaction is not known. But because TA is made up of many galloyl moieties, which each contain multiple vicinal -OH groups, each TA molecule may be able to participate in multiple inhibitory interactions with one or more αS monomers.

It is also likely that TA is not completely oxidized in our samples, and instead may be a mixture of various galloyl and o-quinone intermediates (Fig. S1c). The lysine cross peaks in our NMR spectra are severely overlapped, which does not easily allow for the deconvolution of variable interactions across the αS sequence, and our NMR experiments do not report on changes to the lysine sidechain. However, our data suggests that TA interacts with other residues besides lysine, with a preference for residues with polar sidechains. In particular, the regions of the NAC domain that show peak intensity loss or changes in ^15^N-R_2_ are centered around threonine residues (Fig. 3c). This suggests that the polar residues on αS could be participating in hydrogen bonding interactions with TA galloyl or o-quinone moieties, or that TA stabilizes a compact form of αS where the threonine residues can participate in hydrogen bonding with other residues in αS. This stabilizing behavior was recently identified for the polyphenolic compound Brazilin, which was found to preferentially bind to and stabilize compact conformations of αS [44]. It is also possible that the lysine residues of αS do not conjugate to TA, but rather form stabilizing cation-pi interactions [45] between the positively charged lysine amino sidechain and the aromatic pi-face of the galloyl moieties in TA.

Our NMR data indicate the important role that polar and positively charged amino acids play in interacting with TA. What we observe across the entirety of our NMR data is that TA clearly interacts with a majority of the residues in the N-terminus, rigidifying the residues within this region. The residues that TA appears to interact with in the NAC region and C-terminus are almost exclusively polar or positively charged amino acids, or directly adjacent to these residues. NMR data, along with our ThT assays, AFM, SDS-PAGE, and SEC data indicate that TA interacts with αS and decreases the dynamics along these regions, likely inhibiting the structural transition to a ß-sheet necessary for fibril formation, and forms an assortment of various soluble and insoluble non-fibrillar aggregates.

The N-terminus of αS has recently been found to mediate the interaction between the IDP monomers and the disordered regions of the fibril that initiates the amyloid seeding process [46,47]. The interaction with the disordered regions of amyloid fibrils that make up their “fuzzy” surface is increasingly being identified to be important in more and more amyloid systems [48]. The N-terminal domain has been identified to play a critical role at the start of αS aggregation [15,49], including the identification of a sequence of 7 residues (36-GVLYVGS-42) that was found to play a critical role in the ability of monomeric αS to aggregate [50]. This 7-residue sequence of the N-terminal domain shows the largest increase in ^15^N-R_2_ observed in this work (Fig. 3a,b), suggesting that it may be particularly susceptible to interaction with TA and other polyphenols. The results in this work illustrate the effectiveness in targeting the αS N-terminal domain by a large polyphenol, where it interacts with polar and positively charged residues and inhibits fibril formation. Although TA itself may produce toxic products after undergoing autooxidation at neutral pH [51], and therefore may not represent a good candidate for potential therapies in humans, the chemical moieties of TA can be utilized for design of future therapeutics. There may be other avenues to utilize TA for its potent anti-aggregation activity in combination with other free radical scavenger and antioxidant molecules, as was demonstrated by our previous report [30]. Regardless of the modality used, it is clear that targeting the N-terminus of αS presents promising inhibitory potential for the development of aggregation inhibitors.

## 5. Materials and Methods

### Protein expression and purification

αS plasmids were transformed in *E. Coli* BL21(DE3) cells while expression was carried out in LB media for non-labeled protein and M9 minimal media containing ^15^NH_4_Cl (Cambridge Isotope Laboratories, Tewksbury, MA) for ^15^N. All expression and purification of both labeled and unlabeled αSyn were carried out as explained previously [52]. The weight and purity of αSyn was confirmed via SDS-PAGE and HSQC.

### NMR sample preparation and experiments

Lyophilized ^15^N-labeled protein powder was dissolved in buffer (20 mM sodium phosphate, 100 mM NaCl). The protein was then filtered through a 100-kDa filter to remove any higher-ordered aggregates and then concentrated using a 3-kDa filter (Millipore Sigma, St. Louis, MO). Samples were diluted to a final protein concentration of 150-300 μM with 10% D2O added. ^1^H-^15^N HSQC and If sample contained tannic acid (Sigma Aldrich, St. Louis, MO), pH was checked after addition of tannic acid to ensure no changes in pH. ^1^H-^15^N HSQC and ^15^N-^13^C CON were taken at 150 μM. The αS R_2_ Relaxation experiment was taken at 300 μM and the αS:TA R_2_ Relaxation experiment was conducted at 250 μM. The ^1^H-^15^N HSQC, ^15^N-^13^C CON, and R_2_ Relaxation experiments were recorded on Bruker Avance III spectrometers at either 600 or 700 MHz ^1^H Larmor frequency. Spectra were processed using Bruker Topspin and the data was analyzed using CCPNMR v3.

### Fibril sample preparation

Lyophilized protein was dissolved in 10 mM phosphate-buffered saline (PBS) (pH 7.4). Large aggregates were removed by filtering through a 100-kDa filter followed by concentrating the solution in a 3-kDa filter (Millipore Sigma). To create fibrils, 0.5-1.0 mL of 100 μM of protein are shaken for two weeks at 300 rpm and 37°C in PBS. Samples used for AFM were collected by centrifugation at 14k rpm for 2 hours and washed through resuspension with PBS (pH 7.4). This process is repeated as necessary to remove residual soluble and non-fibrillar components.

### AFM sample preparation

Samples (20 μL) were pipetted onto freshly cleaved mica (Ted Pella Inc., Redding, CA) and incubated for 20 min at room temperature, followed by 3 washes of 150 μL deionized water. If imaging in air, samples were dried for one hour before imaging. All images were collected on a Cypher ES instrument (Asylum Research, Oxford Instruments, Goleta, CA) using AC240TS-R3 non-contact mode tips (Asylum Research, Oxford Instruments). Image processing and analysis were carried out in the IgorPro software package (WaveMetrics, Portland, OR).

### ThT fluorescence assay

Lyophilized protein was dissolved in PBS buffer at the desired pH (6.0 or 7.4). Any large aggregates were removed by filtering the solution through a 100-kDa filter and then concentrating the solution using a 3-kDa filter (Millipore Sigma). Samples were diluted to 70 μM, along with 20 μM ThT (Acros Organics, Pittsburgh, PA) and various concentrations of tannic acid. The samples were then loaded into a 96-well plate (Corning, Corning, NY). A single Teflon bead (3 mm; Saint-Gobain N.A., Malvern, PA) was added to each well and then the plate was then sealed with tape. Plates were shaken at 37°C and 600 rpm until the increase in ThT fluorescence intensity (480 nm), which was measured every 33 min by a POLAR Star Omega plate reader (BMG Labtech, Cary, NC), had plateaued. At least five replicates of the ThT assay were recorded for each sample. Each fluorescence trace is displayed with the averages of the total runs along with the standard deviation present as the error.

### Sodium dodecyl sulphate polyacrylamide gel electrophoresis (SDS-PAGE)

Solutions of 50 μM αSyn (pH 6.0 and 7.4) were prepared via methods previously reported here. The αSyn samples were incubated alone and with various concentrations of tannic acid over the course of a week. After 7 days, the solutions were centrifuged for 1.5 hrs at 14k rpm. After centrifugation, the supernatant was removed and incubated with 8 M guanidine hydrochloride (Thermo Fisher Scientific, Waltham, MA) for 2 hrs to denature any aggregates. The samples were then washed using water and 3-kDa filters (Millipore Sigma) to remove any guanidine hydrochloride. 20 μL of sample was then mixed with a gel loading dye (Thermo Fisher Scientific). The mixture was then loaded into a 15% SDS-PAGE gel. Electrophoresis was performed, 100 V for 10 min and 200 V for 30 min, and the gel was stained with Simply Blue SafeStain (Thermo Fisher Scientific) for 30-60 min. The gel was then de-stained with a solution of 10% acetic acid, 40% ethanol, and 50% distilled water overnight. The gel was imaged with an Aplegen Omega Fluor Gel Documentation System and software (Gel Company Inc., San Francisco, CA). The bands were quantified using the Image J software (NIH, Bethesda, MD).

### Separating aggregates via size-exclusion chromatography (SEC)

Lyophilized protein was dissolved in preparation to form fibrils via the method previously reported in this section. Once the protein solution was ready, tannic acid was added to the solution to achieve the desired ratio of αSyn:tannic acid. The samples were then shaken for a specific time interval at 300 rpm and 37°C in PBS. After 1-24 hrs, 0.5 mL of sample was injected into the sample loop of the Akta Pure fast protein liquid chromatography instrumentation (GE, Boston, MA). The sample was then separated via the Superose 6 Increase 10/300 GL column (GE) and collected via a fractional collector. The buffer used during separation was PBS with a flow rate of 0.250 mL/min.

### Tannic acid high-pressure liquid chromatography (HPLC) and UV-Vis characterization

Tannic acid and gallic acid (Sigma Aldrich) were dissolved in 150 μM of a combination of citric acid (Sigma Aldrich) and sodium phosphate (Sigma Aldrich) buffer, the ratio of both was altered to achieve the desired pH’s of either 6.0 or 7.4. The UV-Visible spectrum of each sample was recorded for both tannic acid and gallic acid for both pH’s using a DU730 UV-Visible Spectrometer (Beckman Coulter, Pasadena, CA). The spectra were recorded from 250-400 nm with 0.2 nm data intervals. The tannic acid and gallic acid solutions were diluted until the wavelength of maximum absorbance had an absorbance between 0.5-1.0 mAu. The HPLC profile of tannic acid and gallic acid at both pH’s was obtained using an Infinity II HPLC system (Agilent Technologies, Santa Clara, CA) and an Agilent 5 prep-C18 column (50×21.2 mm; Agilent Technologies). The observed wavelength was 280 nm for TA and 260 nm for GA. For the HPLC, the citric acid/sodium phosphate buffer was used as the aqueous phase and HPLC-grade acetonitrile (Sigma Aldrich) was used as the organic phase. All buffers were sonicated for 20 min before use in the HPLC to ensure bubbles did not get into HPLC system.

## Supplementary Materials

Supplementary information available online includes Figures S1-S6.

## Data Availability

All data used in this work is included in the main text or supplementary information files. Additional requests for data can be directed to the corresponding author.

## Author Contributions

J.S., P.V.M. and J.B. conceptualized the project and designed the experiments. J.S. performed the experiments and collected data. J.S. and J.K.W. analyzed the data. All authors contributed to writing and editing the manuscript, and approved the final version.

## References

1. Snead, D.; Eliezer, D. Alpha-Synuclein Function and Dysfunction on Cellular Membranes. Experimental Neurobiology 2014, 23, 292–313, doi: 10.5607/en.2014.23.4.292.

2. Jain, N.; Bhasne, K.; Hemaswasthi, M.; Mukhopadhyay, S. Structural and Dynamical Insights into the Membrane-Bound α-Synuclein. PLOS ONE 2013, 8, e83752, doi: 10.1371/journal.pone.0083752.

3. Zhu, M.; Fink, A.L. Lipid Binding Inhibits α-Synuclein Fibril Formation*. Journal of Biological Chemistry 2003, 278, 16873–16877, doi:https://doi.org/10.1074/jbc.M210136200.

4. Bartels, T.; Choi, J.G.; Selkoe, D.J. α-Synuclein occurs physiologically as a helically folded tetramer that resists aggregation. Nature 2011, 477, 107–110, doi: 10.1038/nature10324.

5. Lee, H.J.; Choi, C.; Lee, S.J. Membrane-bound alpha-synuclein has a high aggregation propensity and the ability to seed the aggregation of the cytosolic form. J Biol Chem 2002, 277, 671–678, doi:10.1074/jbc.M107045200.

6. Apetri, M.M.; Maiti, N.C.; Zagorski, M.G.; Carey, P.R.; Anderson, V.E. Secondary Structure of α-Synuclein Oligomers: Characterization by Raman and Atomic Force Microscopy. Journal of Molecular Biology 2006, 355, 63–71, doi: https://doi.org/10.1016/j.jmb.2005.10.071.

7. Roeters, S.J.; Iyer, A.; Pletikapić, G.; Kogan, V.; Subramaniam, V.; Woutersen, S. Evidence for Intramolecular Antiparallel Beta-Sheet Structure in Alpha-Synuclein Fibrils from a Combination of Two-Dimensional Infrared Spectroscopy and Atomic Force Microscopy. Scientific Reports 2017, 7, 41051, doi:10.1038/srep41051.

8. Lorenzen, N.; Lemminger, L.; Pedersen, J.N.; Nielsen, S.B.; Otzen, D.E. The N-terminus of α-synuclein is essential for both monomeric and oligomeric interactions with membranes. FEBS Letters 2014, 588, 497–502, doi: https://doi.org/10.1016/j.febslet.2013.12.015.

9. Vamvaca, K.; Volles, M.J.; Lansbury, P.T. The First N-terminal Amino Acids of α-Synuclein Are Essential for α-Helical Structure Formation In Vitro and Membrane Binding in Yeast. Journal of Molecular Biology 2009, 389, 413–424, doi: https://doi.org/10.1016/j.jmb.2009.03.021.

10. Tuttle, M.D.; Comellas, G.; Nieuwkoop, A.J.; Covell, D.J.; Berthold, D.A.; Kloepper, K.D.; Courtney, J.M.; Kim, J.K.; Barclay, A.M.; Kendall, A.; et al. Solid-state NMR structure of a pathogenic fibril of full-length human α-synuclein. Nature Structural & Molecular Biology 2016, 23, 409–415, doi:10.1038/nsmb.3194.

11. Li, Y.; Zhao, C.; Luo, F.; Liu, Z.; Gui, X.; Luo, Z.; Zhang, X.; Li, D.; Liu, C.; Li, X. Amyloid fibril structure of α-synuclein determined by cryo-electron microscopy. Cell Research 2018, 28, 897–903, doi:10.1038/s41422-018-0075-x.

12. Lautenschläger, J.; Stephens, A.D.; Fusco, G.; Ströhl, F.; Curry, N.; Zacharopoulou, M.; Michel, C.H.; Laine, R.; Nespovitaya, N.; Fantham, M.; et al. C-terminal calcium binding of α-synuclein modulates synaptic vesicle interaction. Nature Communications 2018, 9, 712, doi: 10.1038/s41467-018-03111-4.

13. Kim, T.D.; Paik, S.R.; Yang, C.-H. Structural and Functional Implications of C-Terminal Regions of α-Synuclein. Biochemistry 2002, 41, 13782–13790, doi:10.1021/bi026284c.

14. Dasari, A.K.R.; Kayed, R.; Wi, S.; Lim, K.H. Tau Interacts with the C-Terminal Region of α-Synuclein, Promoting Formation of Toxic Aggregates with Distinct Molecular Conformations. Biochemistry 2019, 58, 2814–2821, doi:10.1021/acs.biochem.9b00215.

15. Williams, J.K.; Yang, X.; Atieh, T.B.; Olson, M.P.; Khare, S.D.; Baum, J. Multi-Pronged Interactions Underlie Inhibition of α-Synuclein Aggregation by β-Synuclein. Journal of Molecular Biology 2018, 430, 2360–2371, doi:https://doi.org/10.1016/j.jmb.2018.05.024.

16. Williams, J.K.; Yang, X.; Baum, J. Interactions between the Intrinsically Disordered Proteins β-Synuclein and α-Synuclein. PROTEOMICS 2018, 18, 1800109, doi: https://doi.org/10.1002/pmic.201800109.

17. Caruana, M.; Högen, T.; Levin, J.; Hillmer, A.; Giese, A.; Vassallo, N. Inhibition and disaggregation of α-synuclein oligomers by natural polyphenolic compounds. FEBS Letters 2011, 585, 1113–1120, doi:https://doi.org/10.1016/j.febslet.2011.03.046.

18. Porat, Y.; Abramowitz, A.; Gazit, E. Inhibition of Amyloid Fibril Formation by Polyphenols: Structural Similarity and Aromatic Interactions as a Common Inhibition Mechanism. Chemical Biology & Drug Design 2006, 67, 27–37, doi:https://doi.org/10.1111/j.1747-0285.2005.00318.x.

19. Dhouafli, Z.; Cuanalo-Contreras, K.; Hayouni, E.A.; Mays, C.E.; Soto, C.; Moreno-Gonzalez, I. Inhibition of protein misfolding and aggregation by natural phenolic compounds. Cellular and Molecular Life Sciences 2018, 75, 3521–3538, doi: 10.1007/s00018-018-2872-2.

20. Mythri, R.B.; Bharath, M.M.S. Curcumin: A Potential Neuroprotective Agent in Parkinson’s Disease. Current Pharmaceutical Design 2012, 18, 91–99, doi:http://dx.doi.org/10.2174/138161212798918995.

21. Bieschke, J.; Russ, J.; Friedrich, R.P.; Ehrnhoefer, D.E.; Wobst, H.; Neugebauer, K.; Wanker, E.E. EGCG remodels mature α-synuclein and amyloid-β fibrils and reduces cellular toxicity. Proceedings of the National Academy of Sciences 2010, 107, 7710, doi: 10.1073/pnas.0910723107.

22. Palazzi, L.; Bruzzone, E.; Bisello, G.; Leri, M.; Stefani, M.; Bucciantini, M.; Polverino de Laureto, P. Oleuropein aglycone stabilizes the monomeric α-synuclein and favours the growth of non-toxic aggregates. Scientific Reports 2018, 8, 8337, doi: 10.1038/s41598-018-26645-5.

23. Meng, X.; Munishkina, L.A.; Fink, A.L.; Uversky, V.N. Molecular Mechanisms Underlying the Flavonoid-Induced Inhibition of α-Synuclein Fibrillation. Biochemistry 2009, 48, 8206–8224, doi:10.1021/bi900506b.

24. Velander, P.; Wu, L.; Hildreth, S.; Vogelaar, N.; Mukhopadhyay, B.; Zhang, S.; Helm, R.; Xu, B. Catechol-Containing Compounds are a Broad Class of Protein Aggregation Inhibitors: Redox State is a Key Determinant of the Inhibitory Activities. bioRxiv 2020, doi: 10.21203/rs.3.rs-122859/v1.

25. Du, W.-J.; Guo, J.-J.; Gao, M.-T.; Hu, S.-Q.; Dong, X.-Y.; Han, Y.-F.; Liu, F.-F.; Jiang, S.; Sun, Y. Brazilin inhibits amyloid β-protein fibrillogenesis, remodels amyloid fibrils and reduces amyloid cytotoxicity. Scientific Reports 2015, 5, 7992, doi:10.1038/srep07992.

26. Teng, Y.; Zhao, J.; Ding, L.; Ding, Y.; Zhou, P. Complex of EGCG with Cu(II) Suppresses Amyloid Aggregation and Cu(II)-Induced Cytotoxicity of α-Synuclein. Molecules 2019, 24, doi: 10.3390/molecules24162940.

27. Xu, Y.; Zhang, Y.; Quan, Z.; Wong, W.; Guo, J.; Zhang, R.; Yang, Q.; Dai, R.; McGeer, P.L.; Qing, H. Epigallocatechin Gallate (EGCG) Inhibits Alpha-Synuclein Aggregation: A Potential Agent for Parkinson’s Disease. Neurochemical Research 2016, 41, 2788–2796, doi: 10.1007/s11064-016-1995-9.

28. Ono, K.; Hasegawa, K.; Naiki, H.; Yamada, M. Anti-amyloidogenic activity of tannic acid and its activity to destabilize Alzheimer’s β-amyloid fibrils in vitro. Biochimica et Biophysica Acta (BBA) - Molecular Basis of Disease 2004, 1690, 193–202, doi: https://doi.org/10.1016/j.bbadis.2004.06.008.

29. Ono, K.; Yamada, M. Antioxidant compounds have potent anti-fibrillogenic and fibril-destabilizing effects for α-synuclein fibrils in vitro. Journal of Neurochemistry 2006, 97, 105–115, doi:https://doi.org/10.1111/j.1471-4159.2006.03707.x.

30. Zhao, N.; Yang, X.; Calvelli, H.R.; Cao, Y.; Francis, N.L.; Chmielowski, R.A.; Joseph, L.B.; Pang, Z.P.; Uhrich, K.E.; Baum, J.; et al. Antioxidant Nanoparticles for Concerted Inhibition of α-Synuclein Fibrillization, and Attenuation of Microglial Intracellular Aggregation and Activation. Frontiers in Bioengineering and Biotechnology 2020, 8, doi: 10.3389/fbioe.2020.00112.

31. Khurana, R.; Coleman, C.; Ionescu-Zanetti, C.; Carter, S.A.; Krishna, V.; Grover, R.K.; Roy, R.; Singh, S. Mechanism of thioflavin T binding to amyloid fibrils. Journal of Structural Biology 2005, 151, 229–238, doi:https://doi.org/10.1016/j.jsb.2005.06.006.

32. Sneideris, T.; Sakalauskas, A.; Sternke-Hoffmann, R.; Peduzzo, A.; Ziaunys, M.; Buell, A.K.; Smirnovas, V. The Environment Is a Key Factor in Determining the Anti-Amyloid Efficacy of EGCG. Biomolecules 2019, 9, doi: 10.3390/biom9120855.

33. Sternke-Hoffmann, R.; Peduzzo, A.; Bolakhrif, N.; Haas, R.; Buell, A.K. The Aggregation Conditions Define Whether EGCG is an Inhibitor or Enhancer of α-Synuclein Amyloid Fibril Formation. International Journal of Molecular Sciences 2020, 21, doi:10.3390/ijms21061995.

34. Katwa, L.C.; Ramakrishna, M.; Rao, M.R.R. Spectrophotometric assay of immobilized tannase. Journal of Biosciences 1981, 3, 135–142, doi:10.1007/BF02702656.

35. Velander, P.; Wu, L.; Ray, W.K.; Helm, R.F.; Xu, B. Amylin Amyloid Inhibition by Flavonoid Baicalein: Key Roles of Its Vicinal Dihydroxyl Groups of the Catechol Moiety. Biochemistry 2016, 55, 4255–4258, doi:10.1021/acs.biochem.6b00578.

36. Eliezer, D.; Kutluay, E.; Bussell, R.; Browne, G. Conformational properties of α-synuclein in its free and lipid-associated states11Edited by P. E. Wright. Journal of Molecular Biology 2001, 307, 1061–1073, doi:https://doi.org/10.1006/jmbi.2001.4538.

37. Kang, L.; Wu, K.-P.; Vendruscolo, M.; Baum, J. The A53T Mutation is Key in Defining the Differences in the Aggregation Kinetics of Human and Mouse α-Synuclein. Journal of the American Chemical Society 2011, 133, 13465–13470, doi:10.1021/ja203979j.

38. Xie, M.; Hansen, A.L.; Yuan, J.; Brüschweiler, R. Residue-Specific Interactions of an Intrinsically Disordered Protein with Silica Nanoparticles and Their Quantitative Prediction. The Journal of Physical Chemistry C 2016, 120, 24463–24468, doi: 10.1021/acs.jpcc.6b08213.

39. Xie, M.; Li, D.-W.; Yuan, J.; Hansen, A.L.; Brüschweiler, R. Quantitative Binding Behavior of Intrinsically Disordered Proteins to Nanoparticle Surfaces at Individual Residue Level. Chemistry – A European Journal 2018, 24, 16997–17001, doi: https://doi.org/10.1002/chem.201804556.

40. Zhu, M.; Rajamani, S.; Kaylor, J.; Han, S.; Zhou, F.; Fink, A.L. The Flavonoid Baicalein Inhibits Fibrillation of α-Synuclein and Disaggregates Existing Fibrils*. Journal of Biological Chemistry 2004, 279, 26846–26857, doi:https://doi.org/10.1074/jbc.M403129200.

41. Masuda, M.; Suzuki, N.; Taniguchi, S.; Oikawa, T.; Nonaka, T.; Iwatsubo, T.; Hisanaga, S.-i.; Goedert, M.; Hasegawa, M. Small Molecule Inhibitors of α-Synuclein Filament Assembly. Biochemistry 2006, 45, 6085–6094, doi:10.1021/bi0600749.

42. Hou, Z.; Sang, S.; You, H.; Lee, M.-J.; Hong, J.; Chin, K.-V.; Yang, C.S. Mechanism of Action of (-)-Epigallocatechin-3-Gallate: Auto-oxidation–Dependent Inactivation of Epidermal Growth Factor Receptor and Direct Effects on Growth Inhibition in Human Esophageal Cancer KYSE 150 Cells. Cancer Research 2005, 65, 8049–8056.

43. Sang, S.; Hou, Z.; Lambert, J.D.; Yang, C.S. Redox properties of tea polyphenols and related biological activities. Antioxid Redox Signal 2005, 7, 1704–1714, doi: 10.1089/ars.2005.7.1704.

44. Nahass, G.R.; Sun, Y.; Xu, Y.; Batchelor, M.; Reilly, M.; Benilova, I.; Kedia, N.; Spehar, K.; Sobott, F.; Sessions, R.B.; et al. Brazilin Removes Toxic Alpha-Synuclein and Seeding Competent Assemblies from Parkinson Brain by Altering Conformational Equilibrium. Journal of Molecular Biology 2021, 433, 166878, doi: https://doi.org/10.1016/j.jmb.2021.166878.

45. Gallivan, J.P.; Dougherty, D.A. Cation-π interactions in structural biology. Proceedings of the National Academy of Sciences 1999, 96, 9459, doi: 10.1073/pnas.96.17.9459.

46. Yang, X.; Wang, B.; Hoop, C.L.; Williams, J.K.; Baum, J. NMR unveils an N-terminal interaction interface on acetylated-α-synuclein monomers for recruitment to fibrils. Proceedings of the National Academy of Sciences 2021, 118, e2017452118, doi: 10.1073/pnas.2017452118.

47. Kumari, P.; Ghosh, D.; Vanas, A.; Fleischmann, Y.; Wiegand, T.; Jeschke, G.; Riek, R.; Eichmann, C. Structural insights into α-synuclein monomer–fibril interactions. Proceedings of the National Academy of Sciences 2021, 118, e2012171118, doi: 10.1073/pnas.2012171118.

48. Ulamec, S.M.; Brockwell, D.J.; Radford, S.E. Looking Beyond the Core: The Role of Flanking Regions in the Aggregation of Amyloidogenic Peptides and Proteins. Frontiers in Neuroscience 2020, 14, doi:10.3389/fnins.2020.611285.

49. Janowska, M.K.; Wu, K.-P.; Baum, J. Unveiling transient protein-protein interactions that modulate inhibition of alpha-synuclein aggregation by beta-synuclein, a pre-synaptic protein that co-localizes with alpha-synuclein. Scientific Reports 2015, 5, 15164, doi:10.1038/srep15164.

50. Doherty, C.P.A.; Ulamec, S.M.; Maya-Martinez, R.; Good, S.C.; Makepeace, J.; Khan, G.N.; van Oosten-Hawle, P.; Radford, S.E.; Brockwell, D.J. A short motif in the N-terminal region of α-synuclein is critical for both aggregation and function. Nature Structural & Molecular Biology 2020, 27, 249–259, doi:10.1038/s41594-020-0384-x.

51. Bolton, J.L.; Trush, M.A.; Penning, T.M.; Dryhurst, G.; Monks, T.J. Role of Quinones in Toxicology. Chemical Research in Toxicology 2000, 13, 135–160, doi:10.1021/tx9902082.

52. Kang, L.; Moriarty, G.M.; Woods, L.A.; Ashcroft, A.E.; Radford, S.E.; Baum, J. N-terminal acetylation of α-synuclein induces increased transient helical propensity and decreased aggregation rates in the intrinsically disordered monomer. Protein Science 2012, 21, 911–917, doi:https://doi.org/10.1002/pro.2088.

