## Supplementary Information for "Tannic Acid Inhibits α-Synuclein Amyloid Fibril Formation via Binding to the Monomer N-terminal Domain"

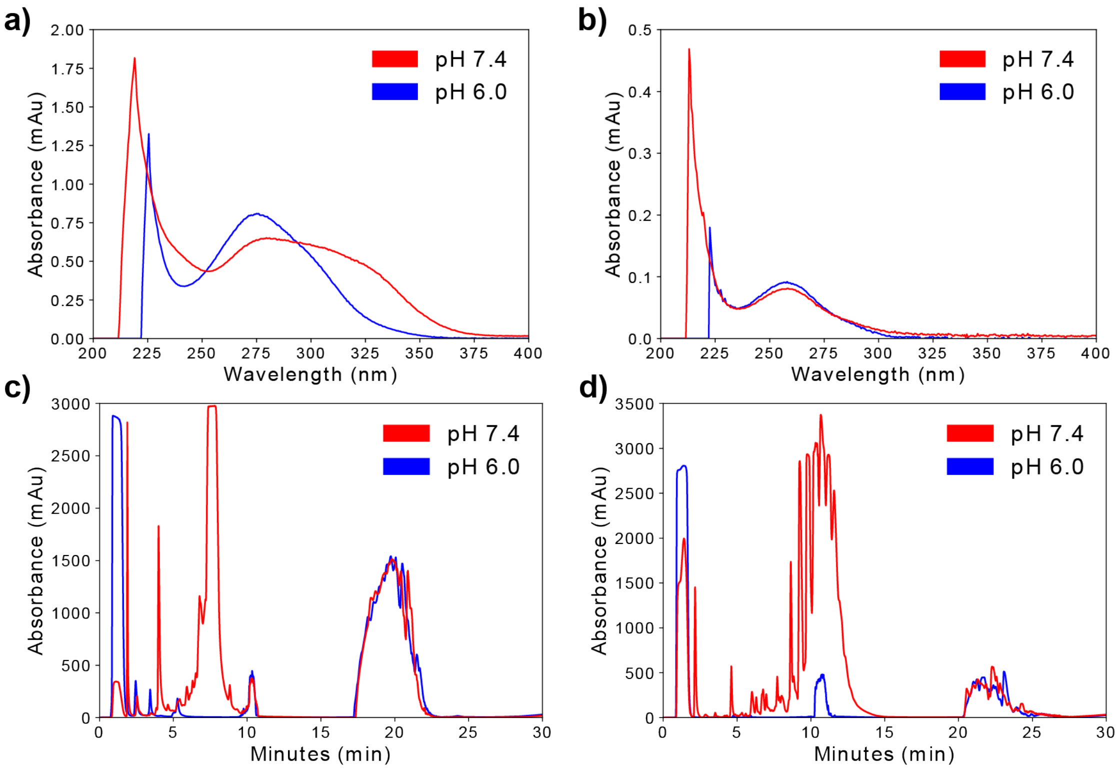


**Figure S1.** UV-Vis spectra of (a) tannic acid and (b) gallic acid, and HPLC elution profiles recorded at (c) 280 nm for tannic acid and at (d) 260 nm for gallic acid. For all four panels (a-d) data collected at pH 7.4 is shown in red, and at pH 6.0 is shown in blue.

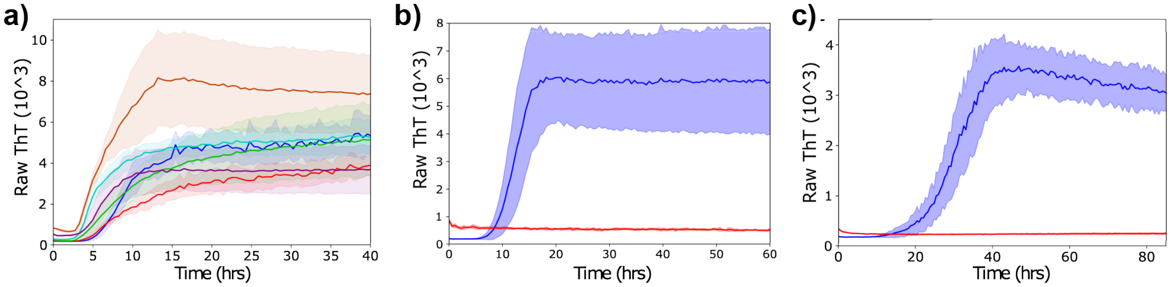


**Figure S2.** (a) ThT fluorescence assays of αS fibril formation with increasing concentrations of TA: control 1:0 (blue), 25:1 (red), 10:1 (green), 5:1 (cyan), 2:1 (purple), 1:1 (orange). All assays were conducted in 100 mM PBS buffer at pH 6.0 with an αS concentration of 70 μM, 37°C and 600 rpm shaking. ThT control assays of 70 μM αS alone (blue) or 70 μM tannic acid alone (red) at (b) pH 6.0 or at (c) pH 7.4.

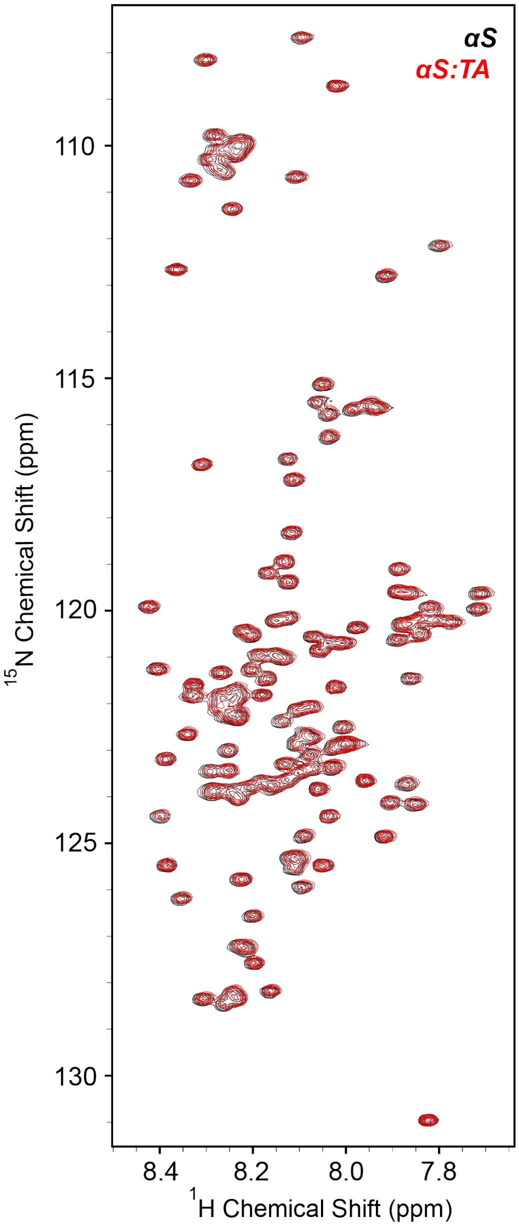


**Figure S3.** $^{1}$H-$^{15}$N HSQC spectra of αS (black) and a 1:1 molar ratio of αS:TA (black) at 4°C in 20 mM sodium phosphate, 100 mM NaCl buffer at pH 6.0.

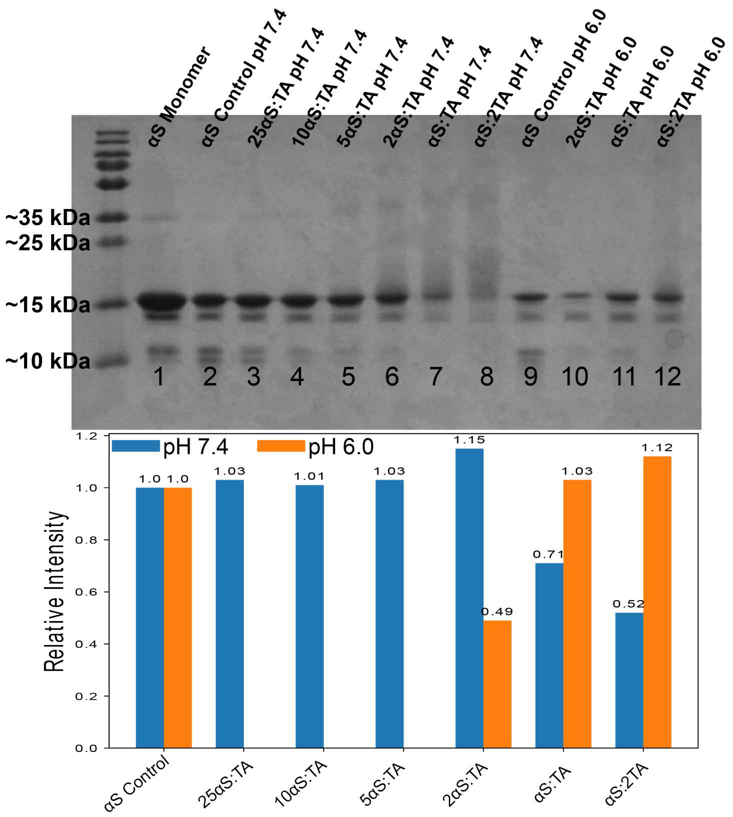


**Figure S4.** SDS-PAGE of the soluble phase of αS with various concentrations of tannic acid after shaking for 5 days at 300 rpm and 37°C, at both pH 7.4 and pH 6.0. The relative intensities for the band at ~15 kDa for each ratio of aS:TA on the gel are shown in the bottom plot, where samples at pH 7.4 are shown in blue and samples at pH 6.0 are shown in orange.

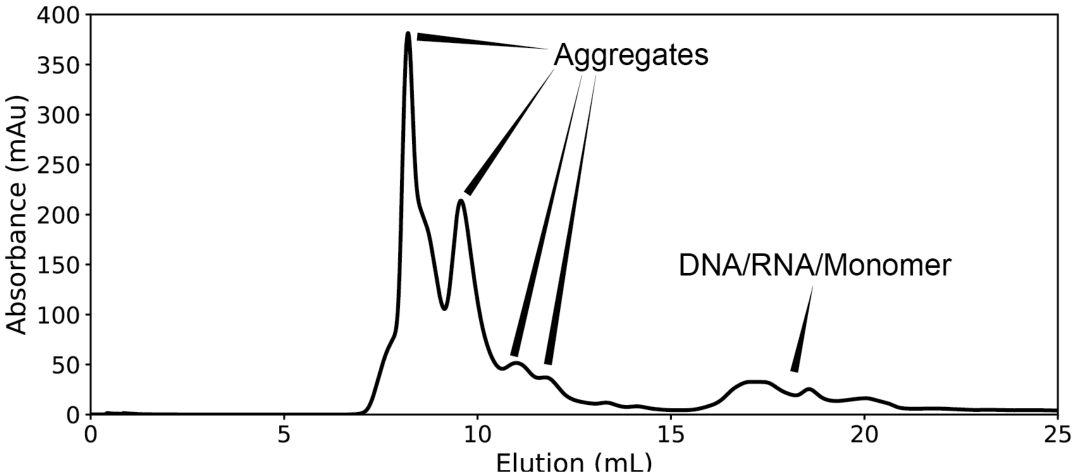


**Figure S5.** SEC elution profile of αS aggregates formed in the presence of tannic acid at a 1:1 molar ratio of αS:TA. The peaks between ~7-12 mL elution volume are the various soluble aggregates formed. The broader peaks around ~16-20 mL elution volume are αS monomers, as well as extraneous DNA/RNA left behind from purification.

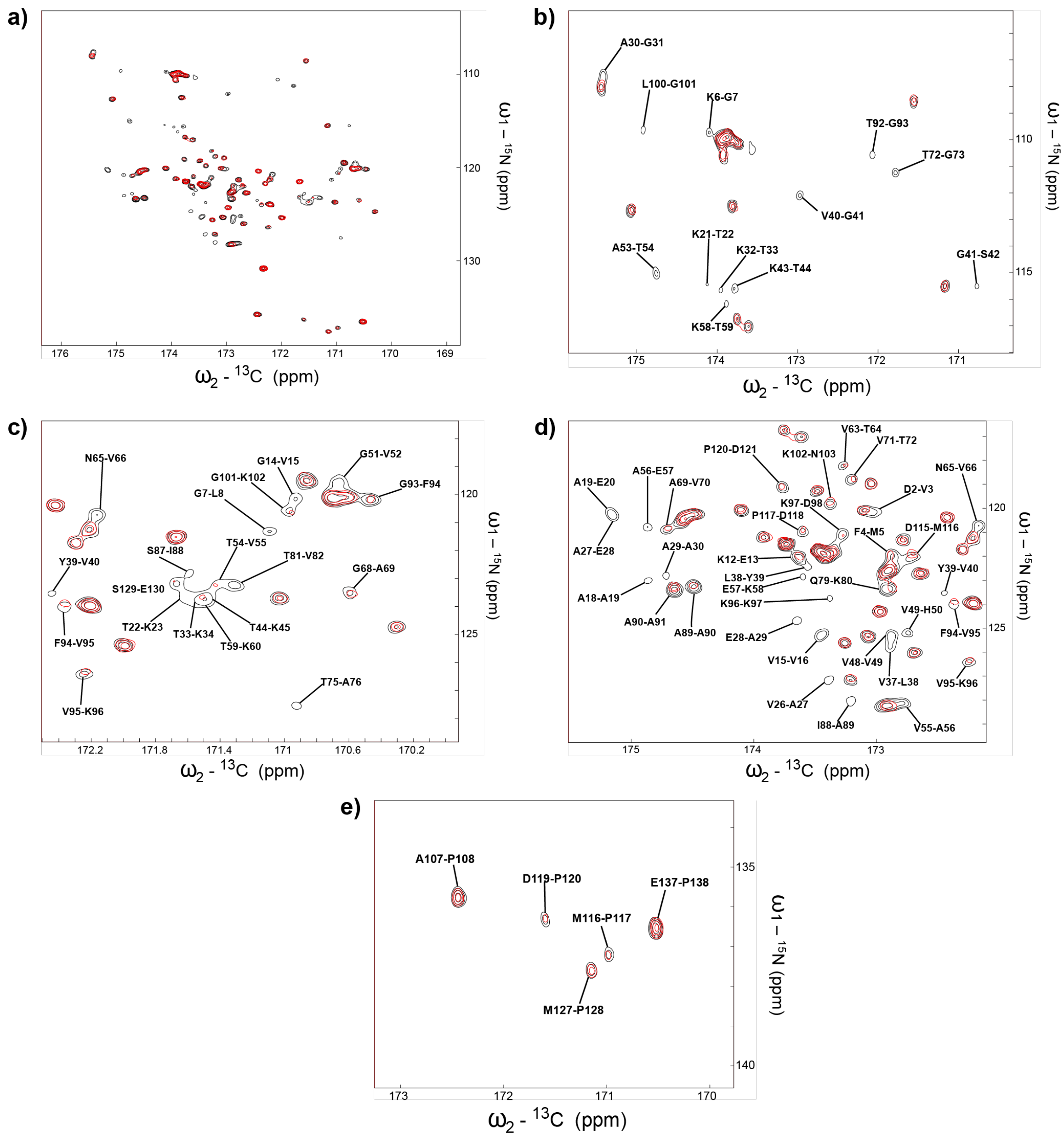

**Figure S5.** $^{15}N$-$^{13}C$ CON spectra of αS (black) and αS:TA at a molar ratio of 1:1 (red). (a) Full spectral window (these are the same spectra shown in Figure 2 of the main text). Panels (b-e) show enlarged parts of the spectrum for ease of viewing.
